# Dissociating the Contributions of Frontal Eye Field Activity to Spatial Working Memory and Motor Preparation

**DOI:** 10.1101/2023.06.12.544653

**Authors:** Donatas Jonikaitis, Behrad Noudoost, Tirin Moore

## Abstract

Neurons within dorsolateral prefrontal cortex of primates are characterized by robust persistent spiking activity exhibited during the delay period of working memory tasks. This includes the frontal eye field (FEF) where nearly half of the neurons are active when spatial locations are held in working memory. Past evidence has established the FEF’s contribution to the planning and triggering of saccadic eye movements as well as to the control of visual spatial attention. However, it remains unclear if persistent delay activity reflects a similar dual role in movement planning and visuospatial working memory. We trained monkeys to alternate between different forms of a spatial working memory task which could dissociate remembered stimulus locations from planned eye movements. We tested the effects of inactivation of FEF sites on behavioral performance in the different tasks. Consistent with previous studies, FEF inactivation impaired the execution of memory-guided saccades, and impaired performance when remembered locations matched the planned eye movement. In contrast, memory performance was largely unaffected when the remembered location was dissociated from the correct eye movement response. Overall, the inactivation effects demonstrated clear deficits on eye movements, regardless of task type, but little or no evidence of a deficit in spatial working memory. Thus, our results indicate that persistent delay activity in the FEF contributes primarily to the preparation of eye movements and not to spatial working memory.

## Introduction

Prefrontal cortex (PFC), particularly dorsolateral PFC, is uniquely evolved among primate species, animals known for their rich behavioral and cognitive repertoire (Miller & Cohen, 2001; Passingham & Wise, 2014). Areas within PFC are highly interconnected not only with one another, but also with both sensory and motor structures, both cortically and subcortically (Preuss & Wise, 2022). Correspondingly, neurons within PFC are known to encode a broad spectrum of signals, including sensory, motor and a variety of cognitive factors. For example, neurons within the macaque frontal eye field (FEF) not only encode the location of visual stimuli and the direction of planned gaze shifts, but also the location of covertly attended (Armstrong et al., 2009; Thompson et al., 2005) or remembered stimulus locations (Armstrong et al., 2009; Bruce & Goldberg, 1985; Hasegawa et al., 2004). Collectively, these properties highlight the conspicuous relationship between the role of PFC in cognition and in its role in premotor processing (Fine & Hayden, 2022; Fuster, 2000; Moore et al., 2003).

The FEF is known to be directly involved in the control of visually guided saccadic eye movements (Bruce & Goldberg, 1985; Dias & Segraves, 1999; Schiller, 1977), and more recently it has been causally implicated in the control of visual spatial attention (Moore & Fallah, 2001, 2004; Thompson et al., 2005). However, its role in spatial working memory remains unclear. Like neurons within other PFC areas (e.g. area 46), many FEF neurons exhibit spatially tuned persistent spiking activity during the delay period of working memory tasks (Armstrong et al., 2009; Clark et al., 2012; Hasegawa et al., 2004). Yet the function of that activity remains ambiguous. A key reason for the ambiguity is that the working memory tasks employed to measure its neural correlates most often do not dissociate the memory and motor demands of the task. In particular, spatial working memory tasks employed in neurophysiological studies in monkeys typically involve delayed movements to remembered locations (Curtis, 2006; Funahashi et al., 1989; Fuster & Alexander, 1971; Wang et al., 2011). Consequently, the possibility of a role of delay activity solely in movement planning, and not working memory, is seldom ruled out. A few studies have employed tasks that deliberately dissociate movement planning from working memory and provide significant evidence that some areas of PFC contribute distinctly to working memory (Funahashi et al., 1993; Hasegawa et al., 2004). However, the function of delay activity within the FEF remains unclear.

Consistent with a role in visually guided saccades and covert spatial attention, local inactivation of neural activity the FEF leads to marked performance deficits in both behaviors (Dias & Segraves, 1999; Monosov & Thompson, 2009). In addition, FEF inactivation is also known to produce sizable performance deficits in a memory-guided saccade (MGS) task (Dias & Segraves, 1999; Noudoost et al., 2014), the task most widely used to elicit persistent delay activity. However, as noted above, such tasks do not dissociate the remembered location from planned movements, as they are identical. Thus, a test of FEF’s contribution to spatial working memory that is distinct from its well-known role in oculomotor programming is needed. We measured the effects of local inactivation of the FEF on the performance of macaque monkeys trained on three different versions of a spatial working memory task. In one version of the task, the location to be remembered was dissociated from the preparation of an eye movement response (Hanning et al., 2016; Hasegawa et al., 2004; Jonikaitis et al., 2019). We observed that FEF inactivation produced clear deficits in saccadic eye movements, regardless of task type. Inactivation also impaired the execution of memory-guided saccades, and impaired performance when remembered locations matched the correct eye movement response. In contrast, memory performance was largely unaffected when the remembered location was dissociated from the correct eye movement response, thus indicating that the FEF contributes primarily to the preparation of eye movements and not to spatial working memory.

## Materials and Methods

### General and Surgical Procedures

Two male rhesus monkeys (Macaca mulatta, 11 and 14 kg), monkey AQ and monkey HB, were used in this study. All surgical and experimental procedures were in accordance with National Institutes of Health Guide for the Care and Use of Laboratory Animals, the Society for Neuroscience Guidelines and Policies, and Stanford University Animal Care and Use Committee. Surgical procedures are detailed in a previous report (Armstrong et al., 2009).

### Behavioral apparatus

Experiments were controlled by a DELL Precision Tower 3620 desktop computer and implemented in Matlab (MathWorks, Natick, MA, USA) using Psychophysics and Eyelink toolboxes (Brainard, 1997; Cornelissen et al., 2002). Eye position was recorded with an SR Research EyeLink 1000 (sampling rate = 1 kHz) desktop mounted eye-tracker for online gaze position tracking and for offline analysis. Stimuli were presented at a viewing distance of 60 cm, on an VIEWPixx3D display (1920 × 1080 pixels, 60 Hz).

### Behavioral tasks

Monkeys were seated in a primate chair in front of the visual display. Monkeys were trained to perform three different versions of a memory task, a memory guided saccade (MGS) task, a ‘Look’ and an ‘Avoid’ task (Hasegawa et al., 2004; Jonikaitis et al., 2019); the latter two are described below. Monkeys initiated behavioral trials by fixating a central fixation spot (blue circle of radius 0.5º visual angle, luminance: 3.8 cd/m^2^, RGB color: 0.08, 0.08, 0.78, with color specified as black: 0, 0, 0 and white: 1, 1, 1), presented on a uniform gray background (10.7 cd/m^2^). After the monkey maintained fixation for 600-800 ms (duration selected randomly on each trial), a cue appeared (colored square frame, size 1º x 1º visual angle) for 50 ms at 4 or 8 randomly selected locations (5º to 7º eccentricity on different sessions). Cue presentation was followed by a delay period that varied randomly from 1400-1600 ms. After the delay period, the fixation spot disappeared, and one of four behavioral response options was possible depending on the specific task (see below). Monkeys received a juice reward for making a correct saccadic eye movement and then maintaining fixation for 200 ms. The intertrial interval was 100 ms after each correct response. Failures to acquire fixation, breaks of fixation during the trial, or incorrect eye movements were not rewarded and were followed by a 2000 ms intertrial interval. All stimuli and task parameters in each new trial were selected randomly and independent from the previous trial.

### Look and Memory guided saccade tasks

During each session, monkeys alternated between two block types. In one of the blocks, monkeys performed the Look task together with randomly interleaved memory guided saccade (MGS) trials (Fig. 1A). In these blocks, the cue color was a black open square for monkey AQ (0.2 cd/m^2^) and a green open square for monkey HB (20.1 cd/m^2^, RGB color: 0.08, 0.78, 0.08). After the delay period on Look trials (∼44% of trials), the fixation spot disappeared and two targets appeared (filled blue circles, radius 1 dva). One of the targets always appeared at the previously cued (matched) location, while the other appeared at one of the other of the 3 (or 7) remaining, randomly selected, locations. Monkeys were rewarded for making a saccadic eye movement to the target at the (matching) cued location. After the fixation spot disappeared on MGS trials (∼44% of trials), monkeys were rewarded making an eye movement to the memorized location of the cue, as in previous studies (Funahashi et al., 1989; Gnadt & Andersen, 1988; Lawrence et al., 2005; Noudoost et al., 2021). On MGS trials, if the saccade landed within 5º visual angle of the cued location, a target appeared (filled blue circle) to confirm a correct response, and reward was delivered.

**Figure 1.**
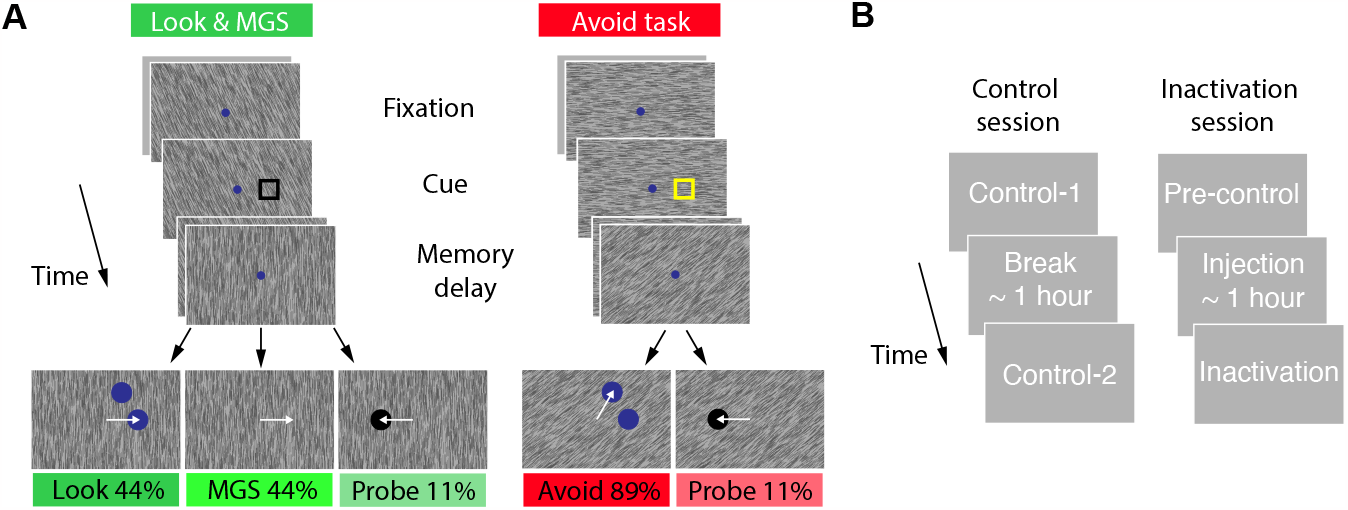
Behavioral task and experimental design. **(A)** Look and Avoid tasks were performed in alternating blocks. In all tasks, a brief visual cue (open square) indicated the location to be memorized. In the Look task (left), after a memory delay, monkeys were rewarded for making a saccade to the target appearing at the cued (matching) location. Memory-guided saccade (MGS) trials were randomly interleaved with Look trials, such that after the memory delay, monkeys were rewarded for making a memory-guided saccade to the cued location. In the Avoid task (right), monkeys made a saccade to the target appearing at the non-cued (non-matching) location. The cue color differed between the Look and Avoid task blocks to highlight the current rule. Randomly interleaved among the Look/MGS and Avoid blocks were a small number of probe trials. On probe trials, moneys were rewarded for making saccades to a single visual target, regardless of where it appeared. **(B)** Control and Inactivation trials were completed both on separate days and during inactivation sessions. During control session, monkeys completed a short block of Look/MGS and Avoid trials (*Control-1*). After a break, monkeys completed a longer block of Look/MGS and Avoid tasks (*Control-2*). During inactivation sessions, monkeys completed a short block of Look/MGS and Avoid trials control trials (*Pre-control-1*). Following FEF inactivation, monkeys completed additional blocks of Look/MGS and Avoid trials.

### Avoid task

In other blocks, monkeys performed the Avoid task. Avoid trials were identical to the Look trials, except that to be rewarded, monkeys made saccadic eye movements to the non-matching target after the delay period (Fig 1A). In this task, the cue color instructed the monkey as to the correct response; green for monkey AQ, black for monkey HB. Each experimental session could begin either with a Look or an Avoid block.

### Probe task

In both block types (Look-MGS and Avoid), probe trials were included in which monkeys were rewarded for making saccades to a single target. Probe trials occurred on ∼11% of total trials and were randomly interleaved with Look-MGS or Avoid trials. In these trials, after the delay, only one target appeared (filled black circle), either at the cued or a non-cued location, and monkeys were rewarded for making a saccadic eye movement to the probe target. These trials are not discussed in this article.

### Control and Inactivation blocks

We collected data during two types of experimental sessions: control and inactivation sessions (Fig. 1B). Control sessions were intended to establish a behavioral baseline and to measure stability of behavior during an experimental session. Inactivation sessions measured the impact of the temporary loss of FEF activity on behavior. Previous studies have shown that the effects of muscimol inactivation typically persist many hours, and well beyond the duration of a single experimental session (Dias & Segraves, 1999), thus, control and inactivation data collection was carried out on separate days. Control sessions were designed to eliminate any confounding effects of time within a session on performance measures. During control sessions, monkeys first completed a short Look-MGS and Avoid task block (*control-1*, 50-100 trials per task) and after a break of ∼1 hour, they completed further blocks of Look-MGS and Avoid tasks (*control-2*, typically ∼250 trials per task based on monkey motivation). The order of Look-MGS and Avoid blocks was randomized in a session. During inactivation sessions, monkeys also first completed a short control block of Look-MGS and Avoid tasks (*pre-control*, 100-200 trials). These data were combined with data from control sessions (*control-1* and *control-2*). Next, after the muscimol infusion (∼1 hour), they completed post-inactivation blocks of Look-MGS and Avoid tasks (*inactivation*).

### FEF inactivation

Prior to inactivation, we located the FEF based on its neurophysiological characteristics and our ability to evoke saccades with electrical stimulation. Electrical microstimulation consisted of 100-ms trains of biphasic current pulses (0.25 ms, 200 Hz) delivered with a Grass stimulator (S88) and two Grass stimulation isolation units (PSIU-6) (Grass Instruments). The FEF was defined as the region from which saccades could be evoked with currents <50 μA (Bruce et al., 1985). In addition, we used 32-channel linear array electrodes with contacts spaced 75 or 150 mm apart (U-Probes and V-Probes, Plexon, Inc) to map out visual and movement-related response fields to corroborate the metrics of the stimulation-evoked saccades (Bruce et al., 1985). Placement of the memory cue during a given inactivation session was determined by the location of evoked saccades and FEF response fields measured on the previous recording/stimulation session at the same FEF site.

During inactivation sessions, we pharmacologically inactivated the FEF via infusion of 0.5-1μL of the GABA_a_ agonist muscimol (5mg/ml), using a custom-made injection system as described and demonstrated previously (Noudoost et al., 2014; Noudoost & Moore, 2011). We did not perform inactivation experiments on consecutive days to provide sufficient recovery from the previous inactivation. Typical duration of the injection was 20 minutes plus an additional period (∼40 minutes) to maximize the expected behavioral effects (Dias & Segraves, 1999; Noudoost et al., 2014).

### Eye movement analysis

Gaze position on each trial was offline drift corrected by using median gaze position from 10 previous trials. Drift correction was based on gaze position from 100 ms to 10 ms before the cue onset, when stable fixation was maintained. We detected saccades offline using an algorithm based on eye velocity changes (Engbert & Kliegl, 2003). We next clustered saccades as ending on one of the three potential locations: (1) fixation, (2) correct response target, (3) wrong response target. The clustering procedure used a support vector machine algorithm with a Gaussian kernel (Jonikaitis et al., 2019). Saccades directed to the target or distractor were required to have a latency of at least 50 ms after the response cue (Fischer & Boch, 1983), and saccades occurring at shorter latency or during the memory delay were classified as fixation breaks. We removed trials if blinks occurred from 100 ms before cue onset to 200 ms after the time of saccade target onset.

### Inactivation effects on performance

For statistical comparisons of paired-means, we drew (with replacement) 10,000 bootstrap samples from the original pair of compared values. We then calculated the difference of these bootstrapped samples and derived two-tailed *p* values from the distribution of these differences. For repeated measures analyses with multiple levels of comparisons, we used one-way and two-way repeated measures ANOVAS. All post-hoc comparisons were based on bootstrap tests and were Bonferonni corrected.

## Results

### Performance during control sessions

Monkeys completed 34 control experiments (AQ n = 15; HB n = 19) and 18 inactivation experiments (AQ n = 15; HB n = 3). Both monkeys performed well in all tasks during control trials. We observed no differences between *Control-1* and *Control-2* blocks during control sessions (p>0.05), and thus combined data from the two along with data from control trials during inactivation sessions (*Pre-control*). In the Look task, mean performance was 95.6±0.7% for monkey AQ and 85.8±1% for monkey HB. In the MGS task, mean performance was 94.6±0.8% for monkey AQ and 87.6±1% for monkey HB. In the Avoid task, mean performance was 88.2±0.5% for monkey AQ and 79.1±2.2% for monkey HB. Performance in the Avoid task was lower than in Look task (difference 8% for monkey AQ and 6.7% for monkey HB, p<0.01), in line with studies that have used multi-task designs, such as pro-saccades and anti-saccades, in the same experimental session (Takeda & Funahashi, 2002)

We also measured saccadic reaction times (RTs) in each of the three tasks. As with performance, we found no differences in RTs between *Control-1* and *Control-2* blocks (ANOVA, p>0.05), and thus combined data from the two along with data from control trials during inactivation sessions (*Pre-control*). Mean RTs during the Look task were 172.4±1.4 ms for monkey AQ and 179.4±0.9 for monkey HB. Reaction times in the MGS task averaged 188.1±2 ms for monkey AQ and 212.5±1.6 ms for monkey HB. In the Avoid task, mean RTs were 161.5±1.4 ms for monkey AQ and 190.6±1.2 ms for monkey HB.

### Effects of inactivation

Figure 2 shows the effects of FEF inactivation on behavioral performance in the 3 different tasks in one session. The effects of inactivation on memory performance are shown along with saccadic RTs in comparison to control data. In the example, the effects of inactivation on behavior during the MGS were most pronounced. Inactivation increased RTs of saccades made into the contralateral (‘inactivated’) hemifield. More significantly, inactivation also reduced the rate of accurate memory-guided saccades made into inactivated hemifield. Both results are consistent with observations from previous studies (Clark et al., 2012; Dias & Segraves, 1999; Sommer & Tehovnik, 1997). In addition, during the Look task, a similar, albeit less pronounced, pair of effects was observed. Saccadic RTs were increased, and performance was reduced in the inactivated hemifield. In contrast, this pattern of results was not observed during the Avoid task. Memory performance in the inactivated hemifield was largely identical to that of the control. In addition, there was no hemifield-specific change in saccadic RTs.

**Figure 2.**
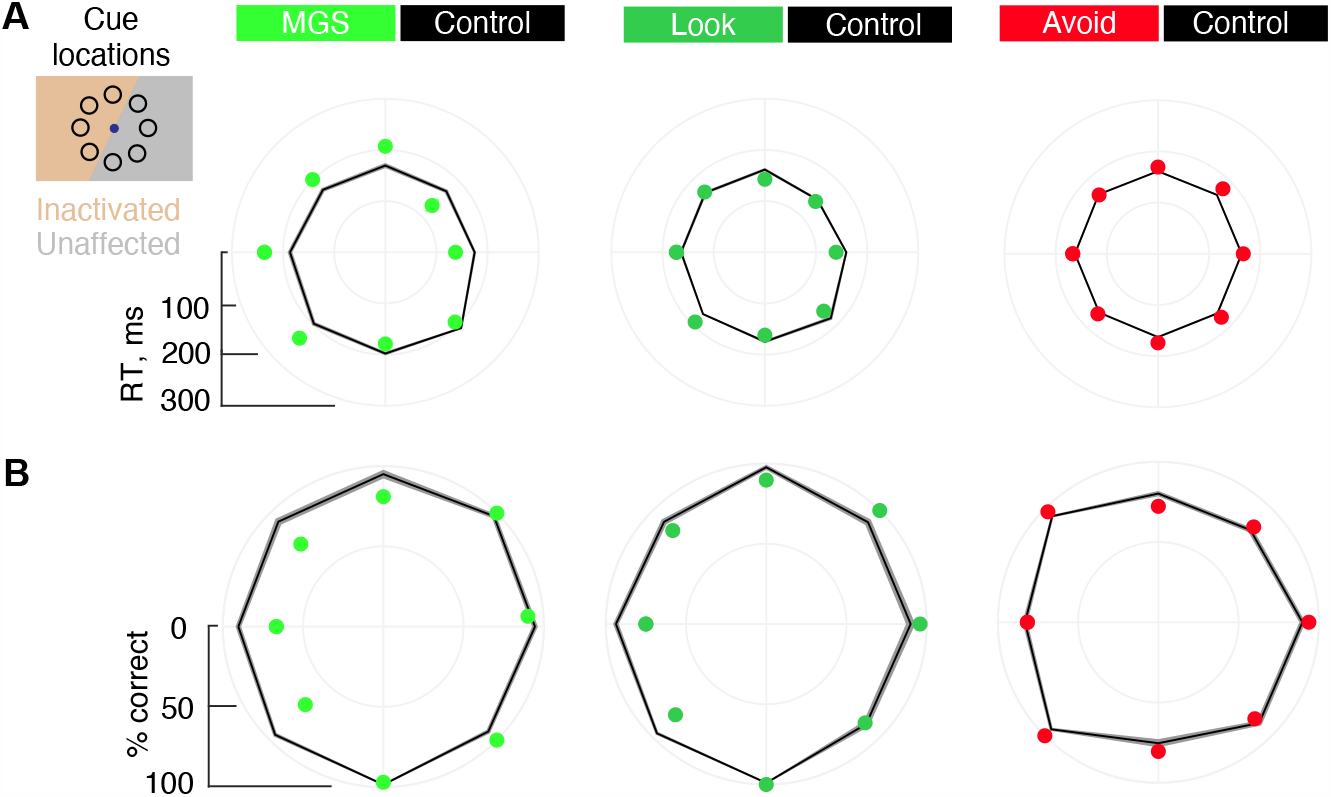
Example inactivation session. Inset shows cue locations used and the division of inactivated and intact visual fields. **(A)** Effects of FEF inactivation on saccadic reaction times. Mean reaction times are shown in polar plots. Colored dots denote saccadic reaction times following FEF inactivation during the MGS, Look and Avoid trials. Black lines indicate the average saccadic reaction times from all control data. **(B)** Effects of FEF inactivation on task performance. Same conventions as in A.

### Effects on saccadic RTs

As in the example session, we focused our analysis on two behavioral measures: saccadic RT and memory performance. We expected to observe clear effects of inactivation on saccadic RT in each form of the memory task based on evidence from previous studies. In contrast the MGS task, saccadic responses in the Look and Avoid tasks were visually guided. Nonetheless, a wealth of past evidence has demonstrated that reversible inactivation of the FEF results in deficits in saccadic RT during both visually guided and memory-guided saccades (Dias & Segraves, 1999; Peel et al., 2017; Sommer & Tehovnik, 1997). Indeed, we observed clear increases in saccadic RTs following FEF inactivation in each form of the memory task (Fig. 3). Differences in RTs compared to control varied significantly as a function of cue location in the MGS task in both monkeys (Fig. 3A) (AQ: F(7)=34.32, *p*<0.0001; HB: F(3)=4.47, *p*=0.014), with increases occurring in the contralateral hemifield. Although smaller in size, the same contralateral RT increases were observed in the Look task in both animals (Fig. 3B) (AQ: F(7)=23.03, *p*<0.0001; HB: F(3)=6.93, *p*=0.0016). In contrast, during the Avoid task, changes in RT did not vary as a function of cue location (Fig 3C) (AQ: F(7)=0.43, *p*=0.88; HB: F(3)=1.48, *p*=0.24). Instead, changes in RT from Control varied significantly as a function of the saccadic target location (AQ: F(7)=11.99, *p*<0.0001; HB: F(3)=4.06, *p*=0.018), with increased RTs during inactivation occurring within the contralateral field (Control - Inactivation difference for contra-lateral hemifield AQ: 23 ms, HB: 9 ms). Thus, when the remembered and movement locations were spatially dissociated, slower RTs were associated with the latter.

**Figure 3.**
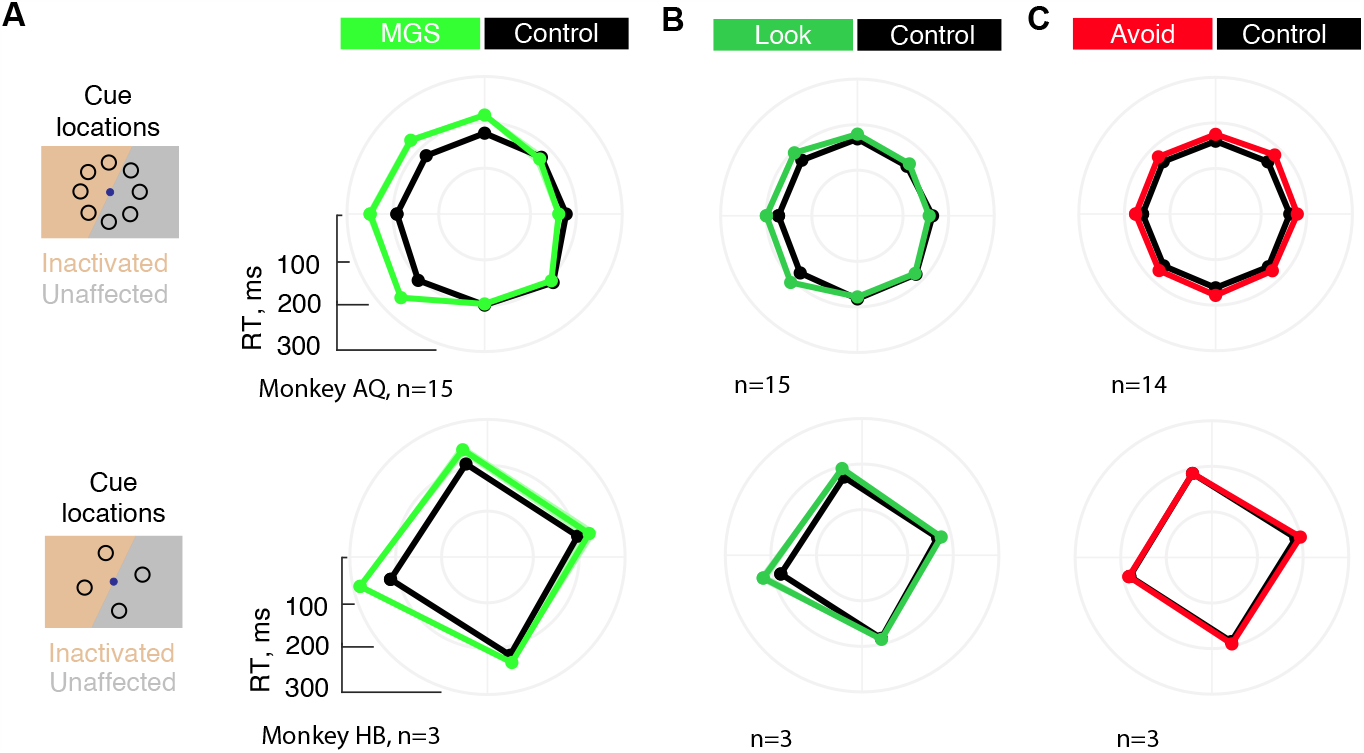
Effects of FEF inactivation on saccadic reaction time across experimental sessions. Data are plotted for monkey AQ (top) and HB (bottom) and for the MGS **(A)**, Look **(B)** and Avoid **(C)** tasks. Shaded error bars denote SEMs.

Next, we examined the effects of inactivation on RTs based on whether the cue and the correct response (saccadic) target appeared within the inactivated or the intact part of visual space (Fig 4). As the two response targets (correct target and distractor) could appear at different (angular) distances from each other, we first examined the effects across different distances between the two targets (Figure 4A). For this analysis, data from the two monkeys were combined. In the Look task, RTs were increased for cues appearing within the inactivated field (F(1)=106.3, *p*<0.0001). Although there was no main effect of distractor distance (F(6)=1.09, *p*=0.36), there was a significant interaction of distractor distance and inactivation (F(6)=3.62, *p*=0.002) in which the smallest distances yielded minimal to no changes in RT. Next, we summarized the data based on whether the cue and/or the distractor appeared in the inactivated or intact field by combining data for different target distances (Fig. 4B). Indeed, during the Look task, increased saccadic RTs were reliably observed within the inactivated visual field regardless of whether or not the distractor also appeared there (*distractor in*: Δ25.6 ± 4.4 ms, *p*<0.001, AQ: Δ26.6 ± 5.1, HB: Δ20.4 ± 6.8; *distractor out*: Δ17.1 ± 2.8 ms, *p*<0.001; AQ: Δ17.2 ± 3.2, HB: Δ16.8 ± 6.4). Saccadic RTs were unaffected when the cue appeared outside of the inactivated field and the distractor appeared within it (*distractor in*: Δ-2.4 ± 2.4 ms, *p*=0.3, AQ: Δ-4.7 ± 2.4, HB: Δ9.3 ± 1.9) and RTs were shorter when both the cue and the distractor appeared outside of the inactivated field (*distractor out*: Δ-11.3 ± 3.7 ms, *p*<0.001; AQ: Δ-13.6 ± 4.3, HB: Δ-0.3 ± 1.5).

**Figure 4.**
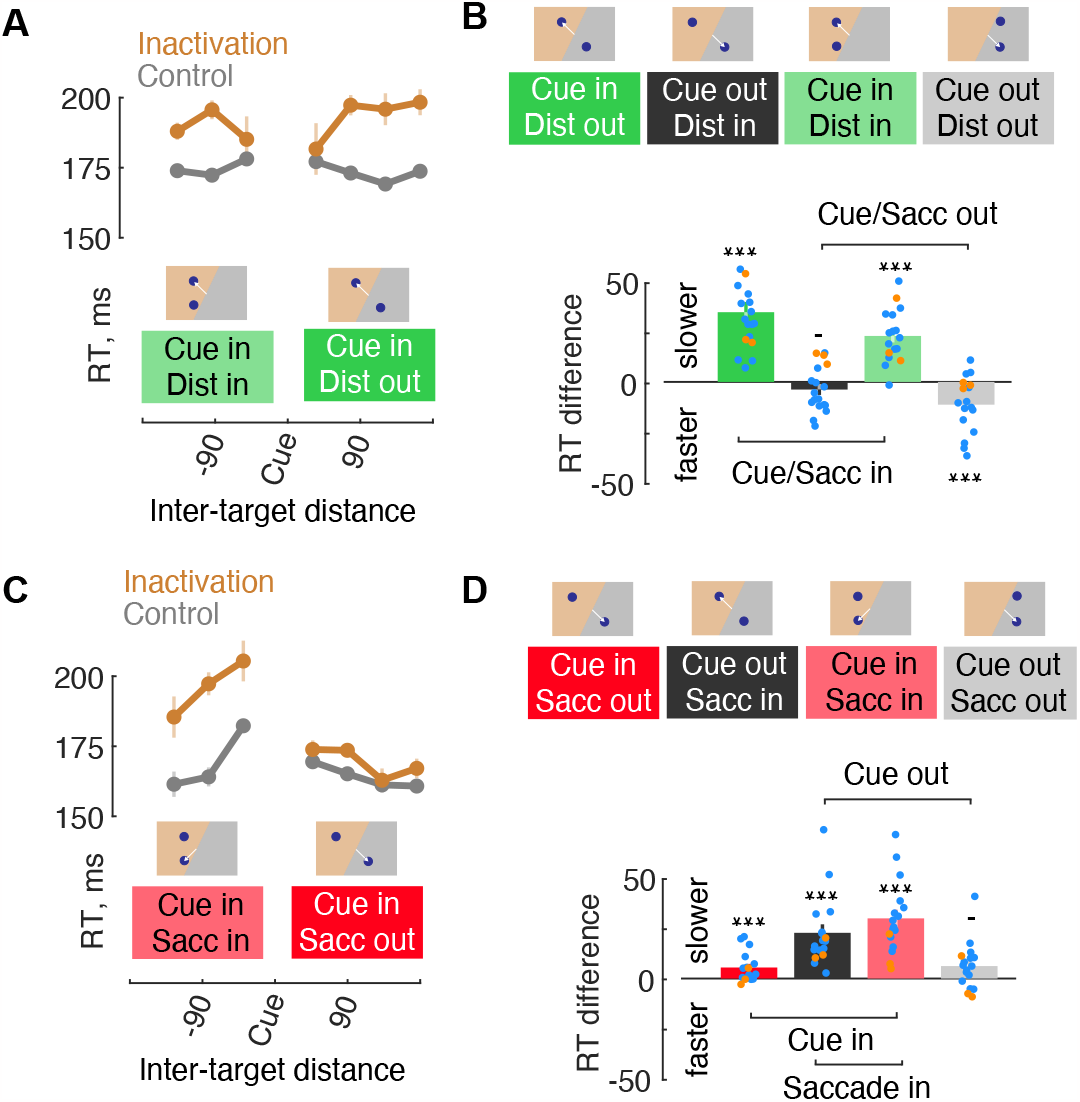
Effects of FEF inactivation on reaction time: cue-distractor and cue-target interactions. **(A)** Reaction times as a function of cue-distractor distance in the Look task. Data are shown for trials in which the cue appeared within the inactivated field and the distractor either also appeared there (Dist in), or appeared in the intact field (Dist out). **(B)** Reaction times for different cue-distractor locations. Top insets illustrate cue and distractor location conditions. White lines indicate the correct saccadic response. Difference in reaction time between inactivation and control data are shown in bar plots (bottom). Dots show individual session data for both monkeys (AQ, blue; HB, orange); error bars denote combined SEMs. **(C)** Same as A, but data are shown from the Avoid task. **(D)** Same as B, but data are shown for the Avoid task.

The pattern of RT effects was different in the Avoid task (Fig 4C). Similar to the Look task, there was a main effect of inactivation (F(1)=53.27, *p*=0.0001). However, there was also significant effect of distance between the cue and saccadic target (F(6)=16.70, *p*<0.0001) in addition to an interaction between distance and inactivation (F(6)=5.92, *p*=0.0001), reflecting the fact that RTs were generally increased for saccades made into the inactivated field. When the data were summarized across target distances based on whether the cue and/or the distractor appeared in the inactivated or intact field, it was clear that increases in RTs largely depending on where the saccade was made (Fig 4D). Saccades made into the inactivated field exhibited large increases in RTs above control regardless of the cue location (*cue out*: Δ20.6 ± 4.2 ms, *p*<0.001; AQ: Δ22.4 ± 5, HB: Δ12 ± 1.7; *cue in*: Δ28.3 ± 4.4 ms, *p*<0.001; AQ: Δ32.6 ± 4.5, ΔHB: 8.7 ± 6.5). In contrast, saccades made into the intact field showed only inconsistent RT effects (*cue out*: Δ5.7 ± 1.9 ms, *p*<0.001; AQ: Δ7 ± 2.1 HB: Δ-0.7 ± 3; *cue in*: Δ3 ± 2.2 ms, *p*=0.17; AQ: Δ4.7 ± 2.2, HB: Δ-5.1 ± 6.2).

### Effects on memory performance

As expected from previous studies, the performance of each monkey on the MGS task decreased in a spatially specific manner following FEF inactivation. Correct saccadic responses to the memorized location decreased contralateral to the FEF site (Fig. 5A). Differences in performance compared to control varied significantly as a function of cue location (F(7)=7.82, *p*<0.0001), with a clear pattern of deficits in the contralateral field. As with saccadic RTs, we observed a less dramatic, but similar pattern of decreased performance in the inactivated field during the Look task (F(7)=5.53, *p*<0.0001). Also as observed with saccadic RTs, the pattern of results was different during the Avoid task. Although differences in performance significantly varied as a function of cue location (F(7)=3.19, *p*=0.003) this effect was due to small decreases in performance observed for cues presented within the intact field, but not the inactivated field.

**Figure 5.**
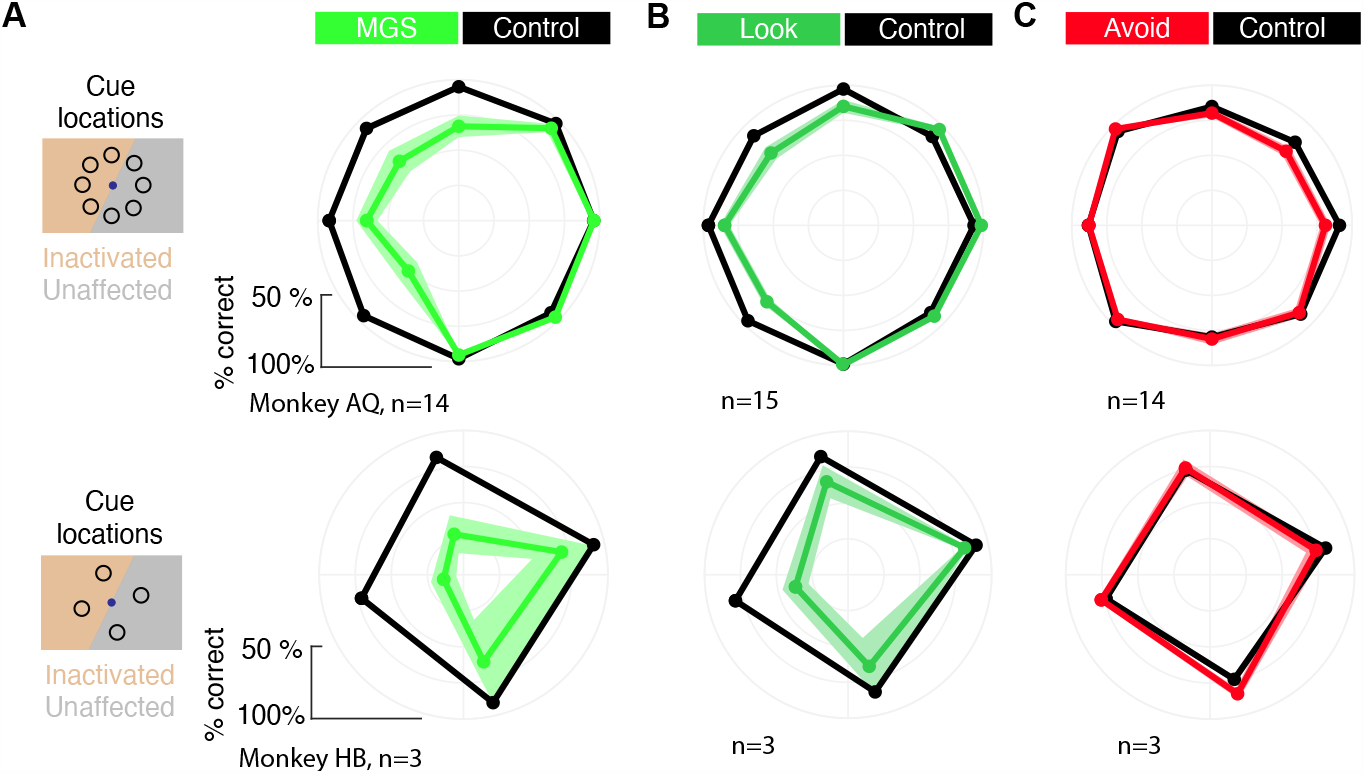
Effects of FEF inactivation on memory task performance across experimental sessions. Data are plotted for monkey AQ (top) and HB (bottom) and for the MGS **(A)**, Look **(B)** and Avoid **(C)** tasks. Shaded error bars denote SEMs.

To further understand the pattern of errors observed in the Avoid task, and further compare it to those in the Look task, we examined the effect of varying the cue and saccadic locations, as in the above RT analyses (Fig 6). In the Look task, inactivation significantly reduced memory task performance (F(1)=71.82, p<0.0001). In addition, there was both a main effect of cue-distractor distance (F(6)=8.64, p= 0.0001) and an interaction between distance and inactivation (F(6)=4.46, p= 0.0001), the latter reflecting the larger deficits when the distractor appeared outside of the inactivated field (Fig. 6A). When the data were summarized across target distances based on whether the cue and/or the distractor appeared in the inactivated or intact field, it was clear that deficits in performance largely depending on where the cue appeared (Fig 6B). Performance in Look task decreased after inactivation when the cue appeared in the inactivated field regardless of where the distractor appeared (*distractor out*: Δ-17.1+3.6%, *p*<0.001, AQ: Δ-13.4 ± 3.3 HB: Δ-35.8 ± 7.1; *distractor in*: Δ-11 ± 2.9%, *p*<0.001; AQ: Δ-10.6 ± 3.4, HB: Δ-13.2 ± 5.7). In contrast, memory performance was not significantly affected when the cue appeared outside of the inactivated field regardless of where the distractor appeared (*distractor in*: Δ-0.1 ± 2.4%, *p*=0.99, AQ: Δ3.1 ± 0.4, HB: Δ-16.1 ± 11.9; *distractor out*: Δ-1.3 ± 3.3%, *p*=0.76; AQ: Δ3.1 ± 1.9, HB: Δ-21.6 ± 11.7). Thus, in the Look task, in which the memorized location and the correct saccadic response were spatially aligned, we observed clear deficits in performance at cued locations within the inactivated field.

**Figure 6.**
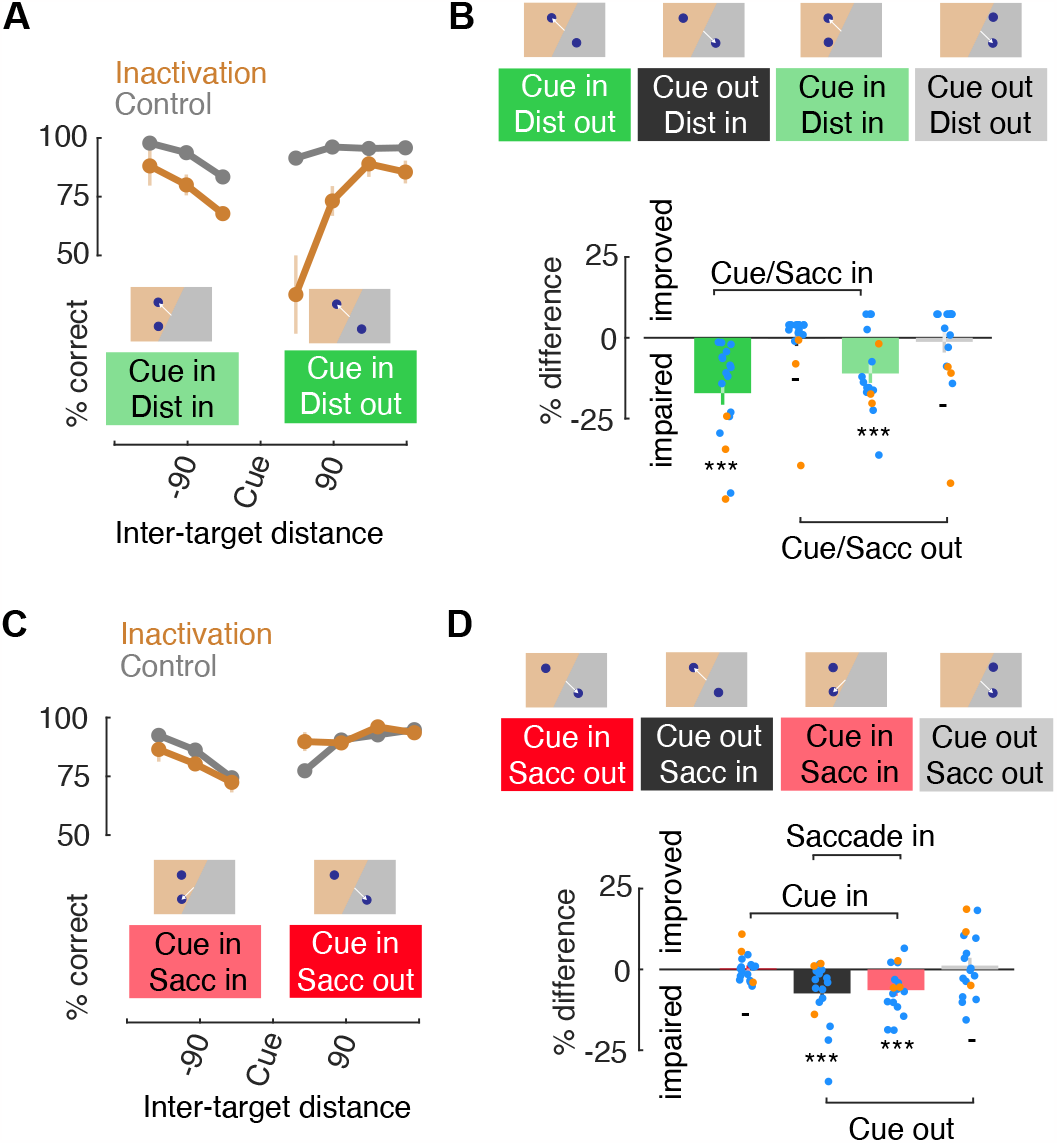
Effects of FEF inactivation on memory task performance: cue-distractor and cue-target interactions. **(A)** performance as a function of cue-distractor distance in the Look task. Data are shown for trials in which the cue appeared within the inactivated field and the distractor either also appeared there (Dist in), or appeared in the intact field (Dist out). **(B)** Performance for different cue-distractor locations. Top insets illustrate cue and distractor location conditions. White lines indicate the correct saccadic response. Difference in reaction time between inactivation and control data are shown in bar plots (bottom). Dots show individual session data for both monkeys (AQ, blue; HB, orange); error bars denote combined SEMs. **(C)** Same as A, but data are shown from the Avoid task. **(D)** Same as B, but data are shown for the Avoid task.

As with saccadic RTs, the pattern of inactivation-induced effects was very different in Avoid task. First, when we examined the effects of inactivation across different distances between the cue and the saccadic target, we found that there was no main effect of inactivation (F(1)=0, p=0.954). There was, however, a main effect of cue-target distance (F(6)=21.73, p= 0.0001), reflecting the drop in overall performance with smaller distances, and an interaction between cue-target distance and inactivation (F(6)=3.25, p=0.005). Second, we combined data across different target distances to examine effects when the cue and/or the saccade target appeared in the inactivated field, we found that the pattern of results demonstrated a dissociation between the memorized and the saccadic target locations (Fig. 6D). When the cue was presented in the inactivated field and the saccadic target appeared outside of it, memory performance was not significantly affected (*cue in, saccade out:* Δ0.3 ± 1%, *p*=0.81; AQ: Δ-0.6 ± 0.8, HB: Δ4.2 ± 4.4), a result that starkly contrasts the result from the same stimulus condition in the Look task (Fig. 6B). However, when the cue appeared outside of the inactivated field, but the saccade target appeared inside of it, performance was slightly impaired (*cue out, saccade in:* Δ-7.5 ± 2.4%, p<0.001, AQ: Δ-8.3 ± 2.7, HB: Δ-3.7 ± 5.2), again in contrast to what we observed in the same stimulus condition in the Look task. Performance was also slightly impaired when both the cue and saccadic target appeared in the inactivated field (*cue in, saccade in:* Δ-6.5 ± 1.8%, p<0.001; AQ: Δ-7.2 ± 2, HB: Δ-2.8 ± 2.8), and there was no effect of inactivation when both the cue and saccadic target appeared in the intact field (*cue out, saccade out:* Δ1.2 ± 2.4%, *p*=0.62; AQ: Δ-0.4 ± 2.4; HB: Δ8.4 ± 7). These results show that when the saccadic response and the cued locations are dissociated, deficits in performance due to FEF inactivation are confined to the location of saccades rather than the location held in memory.

## Discussion

We examined the effects of local inactivation of the FEF on the performance of macaque monkeys trained on three different versions of a spatial working memory task. In one version of the task (Avoid), the location to be remembered was dissociated from the location of planned eye movement responses, whereas in the other two versions, the two locations were matched. We observed that FEF inactivation produced clear deficits in saccadic eye movements in all three versions of the task. Inactivation also impaired the execution of memory-guided saccades, and impaired performance when remembered locations matched the planned eye movement. However, performance was unaffected when the remembered location was dissociated from the correct eye movement response. When the remembered location and the correct eye movement response were spatially coincident, clear deficits in saccadic RTs and memory performance were observed within the inactivated field. In contrast, in the Avoid task, we observed no deficits in memory performance when the remembered locations fell inside of the inactivated field and the correct eye movement response was made within the intact field. Indeed, no deficits in performance or saccadic RTs were observed in the Avoid task unless the eye movement response was made into the inactivated field. These results indicate that FEF inactivation failed to impair spatial working memory, but instead solely produced deficits in the preparation of eye movements. Below, we discuss the implications and potential limitation of these results.

### Premotor functions of FEF delay activity

A recent optogenetic study demonstrated that silencing FEF neuronal activity solely during the delay period of an MGS task significantly disrupted performance (Acker et al., 2016). Notably, performance also depended on normal activity during both the target and response periods. Thus, FEF delay activity appears necessary, but not sufficient, to support performance on the MGS task. In the present study, pharmacological inactivation of the FEF presumably reduced neuronal activity during all epochs of the task, as shown in a previous study (Noudoost et al., 2014). However, in comparison to cue (sensory) driven or motor response-related activity, the unique function, or functions, of delay activity is much more difficult to ascertain.

Persistent, delay-period activity can be observed in cortical areas involved in motor and premotor control (Crammond & Kalaska, 2000; Gnadt & Andersen, 1988; Inagaki et al., 2019), and has thus been interpreted as playing a key role in motor preparation (e.g. (Kaufman et al., 2010; Mountcastle et al., 1975; Snyder et al., 1997; Svoboda & Li, 2018)). In contrast, within areas of dLPFC, where normal function appears to be necessary for performance of tasks requiring short-term memory of recent stimuli or behavioral responses (Bauer & Fuster, 1976; Goldman-Rakic, 1995; Mishkin & Pribram, 1955), delay-period activity is generally interpreted as supporting working memory (Constantinidis et al., 2018). This interpretation has included delay-period activity observed within the FEF (e.g. (Clark et al., 2012; Funahashi et al., 1989)) However, given the preponderance of evidence of FEF’s role in visually guided saccades (Bruce & Goldberg, 1985; Dias & Segraves, 1999; Schiller, 1977) and in visual spatial attention (Moore & Fallah, 2001; Thompson et al., 2005); attributing a specific function to FEF delay activity is particularly challenging. On the one hand, the view of FEF delay activity as principally motor preparatory seems as compelling as that in other premotor structures. On the other hand, psychophysical evidence indicates that attention and working memory, like gaze control and attention, are heavily interdependent, as are their underlying mechanisms (Jonikaitis & Moore, 2019). How then might the present results illuminate the role of FEF delay-period activity?

The present results provide direct evidence that FEF activity, including delay-period activity, is used primarily for motor preparation rather than spatial working memory. The FEF is directly connected with neurons in most areas of extrastriate visual cortex (Schall et al., 1995; Stanton et al., 1995) and appears to serve as an interface between retinotopic visual cortex and other areas of dLPFC (Miller & Cohen, 2001). This unique anatomical position appears to provide a substrate by which neuronal activity signaling planned gaze shifts, whether executed or not, exert attention-related modulations in visual cortical activity (Ekstrom et al., 2008; Gregoriou et al., 2009; Moore & Armstrong, 2003). That is, movement preparatory activity may serve two roles: to shape the metrics of planned gaze shifts via local connections with FEF motor neurons while simultaneously selecting the sensory properties of movement targets via distal connections with visual cortex (Moore et al., 2003). Indeed, a recent study that identified visual cortex-projecting FEF neurons antidromically found that they disproportionately exhibit delay-period activity, and little or no motor burst activity (Merrikhi et al., 2017). Thus, rather than providing a corollary discharge signal directly to visual cortex, which may instead occur via the thalamus (Sommer & Wurtz, 2008), visual cortex-projecting FEF neurons instead convey information about the planning of potential gaze shifts in the form of delay activity.

### Abstract delay signals beyond the FEF

Importantly, a few past studies have also employed tasks that deliberately dissociate movement planning from working memory. In perhaps the most classic study, Funahashi et al. (Funahashi et al., 1993) describe neurons in area 46 that exhibit delay-period activity that is specific to the remembered cue location, and not (or at least to lesser extent) to the planned saccadic response. Another study studied the activity of prefrontal neurons, including the FEF and area 46, during a task similar to that used here and found that although most neurons exhibited delay preferences for either the Look or Avoid conditions, a significant proportion did not; and instead exhibited delay activity that did not differ significantly (Hasegawa et al., 2004). Although neurophysiological studies such as these do not directly address the causal contributions of prefrontal neurons, they nonetheless suggest possible neuronal mechanisms underlying the performance of tasks in which memorized information and behavioral responses are dissociated. Indeed, there is a wealth of neurophysiological evidence indicating a distinct role of dorsolateral prefrontal cortex in more abstract, i.e. non-motor-effector specific, representations (Miller et al., 2002; White & Wise, 1999). Nonetheless, more causal studies will be needed to determine the precise role of those representations in behavior and the relationship between sensorimotor mechanisms and cognition.

## Acknowledgement

We thank D.A. Lopes, S.N. Cital and S. Baker for technical assistance. This work was supported by NIH EY014924 and EY026877

